# Chromosome-level genome assembly of tree sparrow reveals a burst of new genes driven by segmental duplications

**DOI:** 10.1101/2023.02.19.529176

**Authors:** Shengnan Wang, Yingmei Zhang, Yue Shen, Zhaocun Lin, Yuquan Miao, Yanzhu Ji, Gang Song

## Abstract

The creation of new genes is a major force of evolution. Despite as an important mechanism that generated new genes, segmental duplication (SD) has yet to be accurately identified and fully characterized in birds because the repetitive complexity leads to misassignment and misassembly of sequence. In addition, SD may lead to new gene copies, which makes it possible to test the “out of testis” hypothesis which suggests genes are frequently born with testis-specific expression. Using a high-quality chromosome-level assembly, we performed a systematic analysis and presented a comprehensive landscape of SDs in tree sparrow (*Passer montanus*). We detected co-localization of newly expanded genes and long terminal repeat retrotransposons (LTR-RTs), both of which are derived from SDs and enriched in microchromosomes. The newly expanded genes are mostly found in eight families including *C*_*2*_*H*_*2*_*ZNF, OR, PIM, PAK, MROH, HYDIN, HSF* and *ITPRIPL*. The large majority of new members of these eight families have evolved to pseudogenes, whereas there still some new copies preserved transcriptional activity. Among the transcriptionally active new members, new genes from different families with diverse structures and functions shared a similar testis-biased expression pattern, which is consistent with the “out of testis” hypothesis. Through a case analysis of the high-quality genome assembly of tree sparrow, we reveal that the SDs contribute to the formation of new genes. Our study provides a comprehensive understanding of the emergence, expression and fate of duplicated genes and how the SDs might participate in these processes and shape genome evolution.

## Introduction

The origination of new genes is a fundamental question on genome evolution, and gene duplication is one of the most important mechanisms for new gene formation (Ohno 1970; Long et al. 2003; Kaessmann 2010; Ding et al. 2012). Gene duplication can add new copies of genes in the genome, which provide the raw materials for the evolution of novel gene functions and evolutionary adaptation (Crow and Wagner 2006; Magadum et al. 2013). In many cases, the duplicated genes are part of large duplicated chromosomal segments, while the large (>1 kbp) and highly identical (>90%) segment copies in particular chromosomal regions are referred to as segmental duplications (SDs) (Bailey et al. 2001). Owing to their high sequence identity, SDs can promote non-allelic homologous recombination, as a result, they are known as hotspots of chromosomal rearrangement and copy number variation (Bailey et al. 2004; Sharp et al. 2005; Bailey and Eichler 2006; Perry et al. 2006; Liu et al. 2011).

Although critical in genome evolution and plasticity, SDs may be particularly problematic to be characterized at the genomic level because of the inconspicuousness, large size and high sequence similarity, therefore are frequently the last regions of genomes to be sequenced and assembled (Bailey et al. 2001; Vollger et al. 2022). Birds have become one of the most densely sequenced higher-level animal taxa thanks to the Bird 10,000 Genomes (B10K) Project, however, at present, most of the avian genome assemblies are based on the next-generation sequencing (NGS) technology (Zhang et al. 2014; Feng et al. 2020). Due to the short reads produced by NGS, a large number of assemblies of birds are highly fragmented and insufficient for identification of highly duplicated segments. Although SDs have been studied in diverse animal taxa, especially in the primates (Samonte and Eichler 2002; Bailey and Eichler 2006; She et al. 2008), the characterization of SD genomic landscape is relatively limited in birds.

Advances in long-read genome assembly may help to overcome the issue, and the recent generation of a complete telomere-to-telomere (T2T) human genome (T2T-CHM13) successfully demonstrated sequence resolution of complex SDs (Vollger et al. 2022). To enrich our understanding on SDs organization in birds, we generated a chromosome-level genome assembly of tree sparrow (*Passer montanus*), one of the most common passerine species in China, through the combination of long-read HiFi sequencing technology and Hi-C sequencing. Using the high-quality assembly, we identified the SD contents and analyzed its evolutionary process. We found several distinctive characteristics of SDs in the tree sparrow genome. In addition, we further discussed the possible role of SDs and the duplicated genes in genome evolution. This work provides a reference for understanding the SDs organization and the process of new gene formation in birds.

## Results

### Chromosome-level genome assembly of tree sparrow

We sequenced ∼45× HiFi reads from a male tree sparrow collected from LJX and assembled these reads into a 1.28 Gb genome assembly, consisting of 744 contigs with contig N50 length of 54.42 Mb. About 1.16 Gb sequences (91.49% of the total assembly) of the assembled genome were anchored into 36 pseudo-chromosomes with the help of ∼83× Hi-C sequence data (Supplementary Table 1 and 2). Assembly assessment using Benchmarking Universal Single-Copy Orthologs (BUSCO) (Manni et al. 2021) indicated 96.4% avian gene set were present and complete in the assembled genome, confirming the high quality of our assembly (Supplementary Fig. 1). Compared with the previously published Illumina-based assembly of tree sparrow (Qu et al. 2020), our assembly showed great improvement of continuity and completeness (Supplementary Table 2). Subsequent annotation predicted 21,485 protein coding genes covered 94.5% of the complete BUSCO avian gene set (Supplementary Fig. 1).

**Figure 1.**
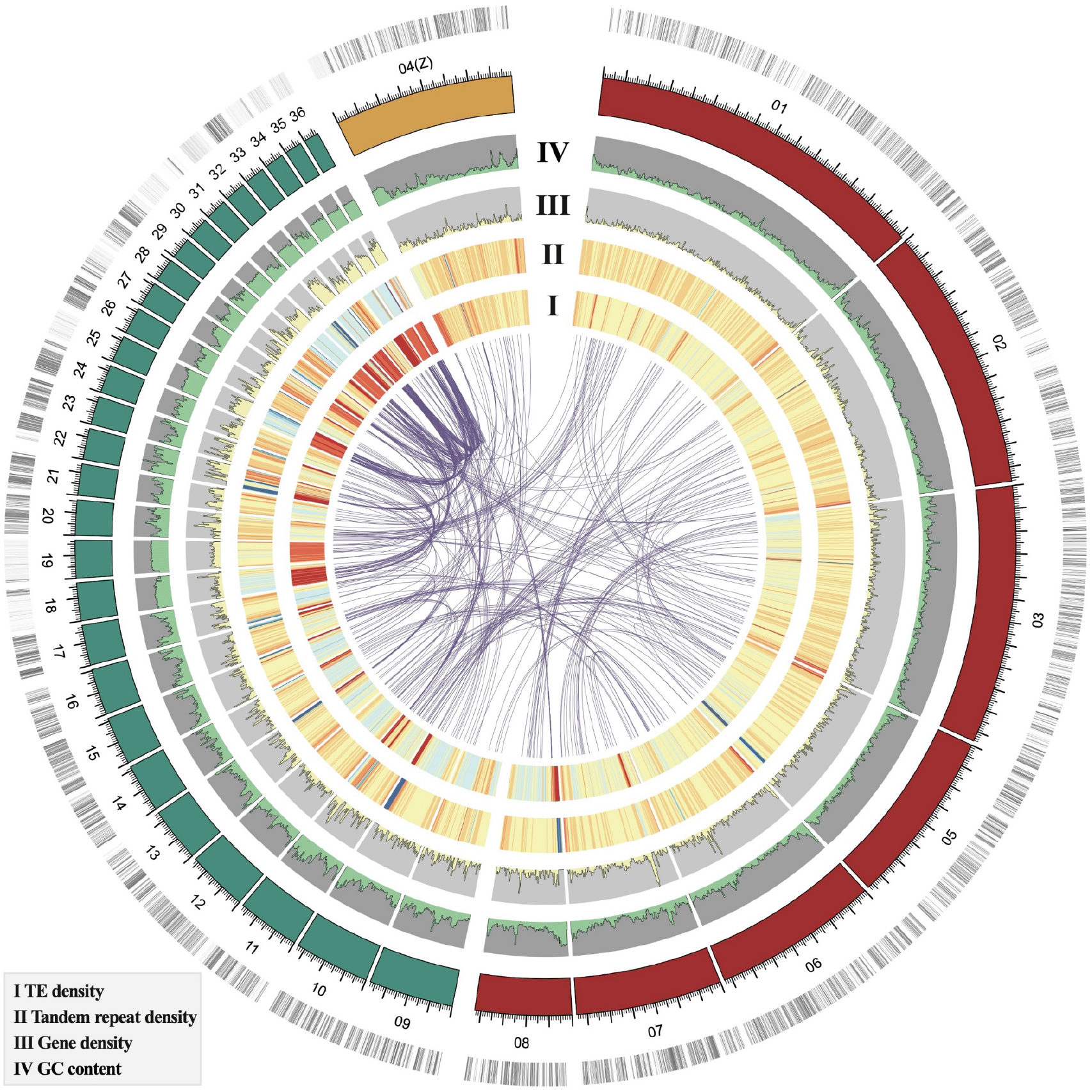
Landscape of assembled tree sparrow genome. The layer of colored blocks is a circular representation of the 36 pseudochromosomes and the outermost track represents the gene distribution in the chromosomes, and we show the microchromosomes in green when the macrochromosomes are shown in red (autosomes) and yellow (Z chromosome). The inner 4 tracks show the GC content, gene density, tandem repeat density and TE density respectively. The synteny blocks are clearly demonstrated by links within the circle.

Tree sparrow has 2n = 78 chromosomes in both sexes, consisting of 8 pairs of relatively large-size macrochromosomes including one pair of sex chromosomes (male ZZ, female ZW), and 31 pairs of smaller microchromosomes (Bulatova et al. 1972). We therefore defined the 8 largest assembled pseudo-chromosomes as macrochromosomes. The macrochromosomes are one-to-one homologous to the large autosomes and chromosome Z of chicken (*Gallus gallus*, GGA), except for chromosomes 2 and 6 which aligned to q-arm and p-arm of GGA1 respectively, and chromosome 5 corresponded to q-arm of GGA4 (Supplementary Table 3 and Supplementary Fig. 2). These exceptions are the results of fission of GGA4 found in different groups of birds, when the fission of GGA1 seems to be apomorphic for Passeriformes (dos Santos et al. 2017; Degrandi et al. 2020). Unlike macrochromosomes, some microchromosomes (chromosomes 18, 19, 25, 27, 30, 31, 32, 34, 35 and 36) showed limited synteny conservation with zebra finch (*Taeniopygia guttata*) and chicken (Supplementary Fig.3).

**Figure 2.**
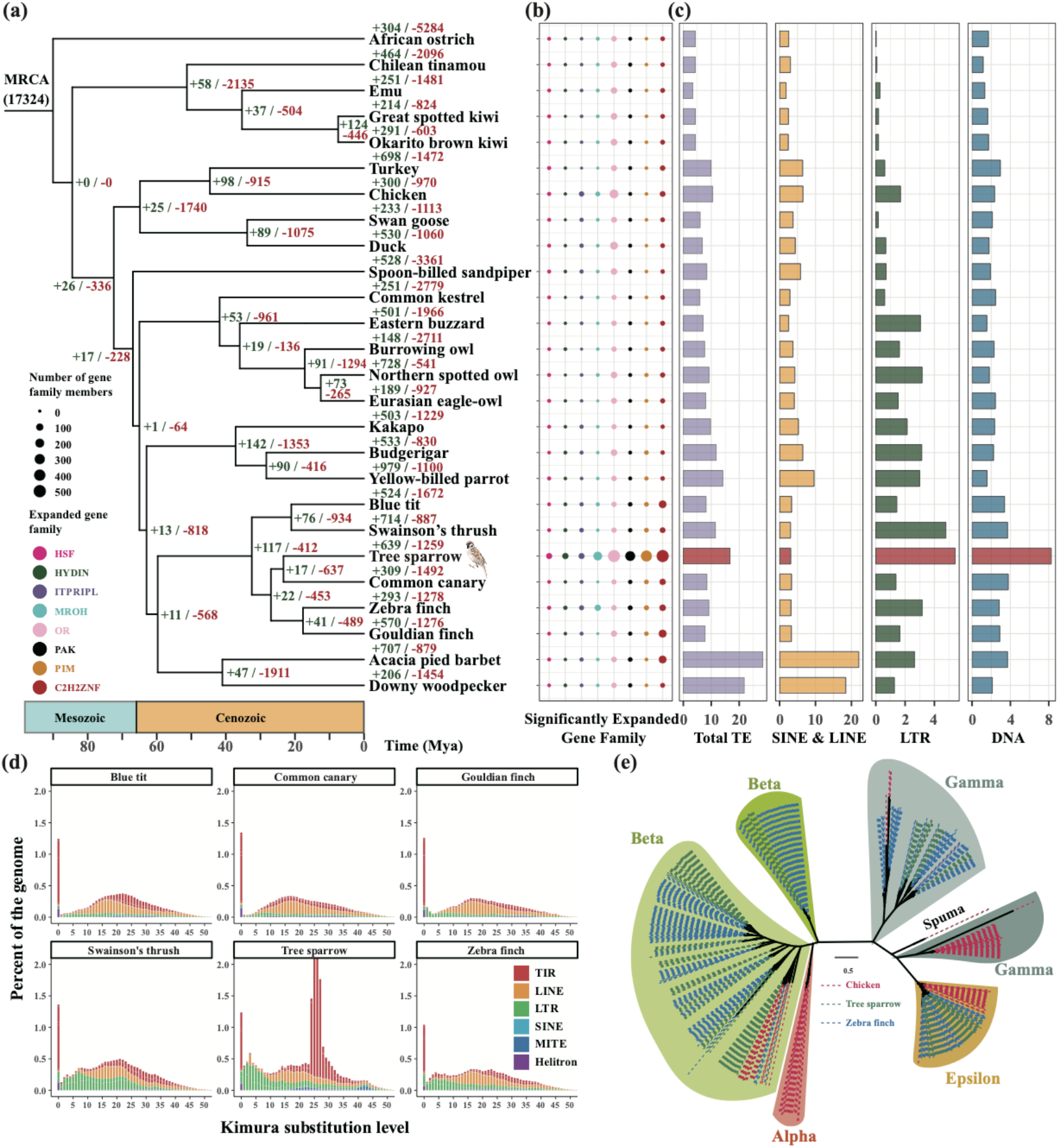
Comparative genomic analysis of tree sparrow. (a) Phylogenetic tree of 26 avian species and gene family evolution. The number of expanded (green) and extracted (red) gene families are shown besides each node and above each species. (b) The eight significantly expanded gene family in tree sparrow. The size of solid circle represents the number of gene family members. (c) Overview of TE contents of 26 avian species. The bar chart displays the percentage of TEs in the assembly. (d) Landscape plot of TE in 6 passerines. Kimura substitution level was showed on the x-axis, and percentage of the genome represented by each TE classification was showed on y-axis. Only the spike at 0% divergence indicating recently active TE. (e) The ML tree of the RT domains of tree sparrow, zebra finch and chicken ERVs.

### Comparative genomics analysis and evolution of gene families

To explore the evolutionary context of tree sparrow, we performed comparative genomic analysis by comparing the tree sparrow genome with another 25 representative avian species (Supplementary Table 6). In total of 4,085 single copy orthologs present in all 26 avian genomes were identified and used to construct a phylogenic tree (Supplementary Table 7). Tree sparrow and common canary (*Serinus canaria*) diverged about 23 million years ago (Mya) (Fig. 2a and Supplementary Fig. 5). The genes in tree sparrow genome were grouped into 13,353 gene families (orthogroups) (Supplementary Fig. 4), among these gene families, 639 expanded and 1,259 contracted (Fig. 2a). In addition, we noticed that there are 8 gene families significantly expanded in tree sparrow, including the Cys_2_His_2_ zinc finger (*C2H2ZNF*) protein, olfactory receptor (*OR*), proviral integration site for Moloney murine leukemia virus (*PIM*), p21-activated kinase (*PAK*), maestro heat-like repeat containing protein family member (*MROH*), hydrocephalus-inducing protein homolog (*HYDIN*), heat shock factor (*HSF*) and inositol 1,4,5-trisphosphate receptor-interacting protein-like (*ITPRIPL*) (Fig. 2b).

**Figure 3.**
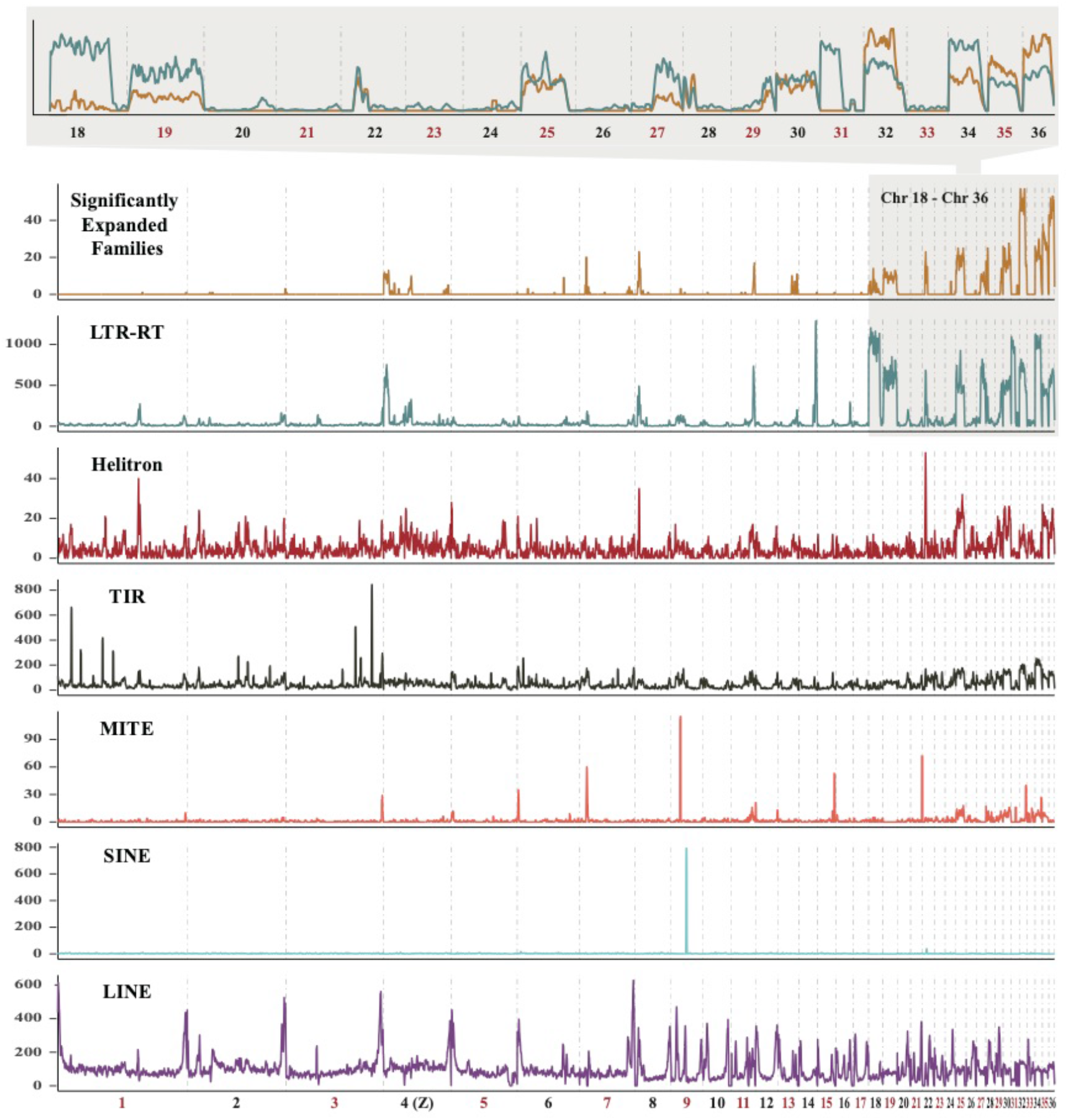
Chromosomal distribution of eight significantly expanded gene families and TEs. The distribution of members of the eight significantly expanded gene families across chromosomes is consistent with the LTR-RTs. The microchromosomes 18-36 are zoomed in, and all members of the eight gene families (yellow) are showed on the uppermost panel with LTR-RTs.

**Figure 4.**
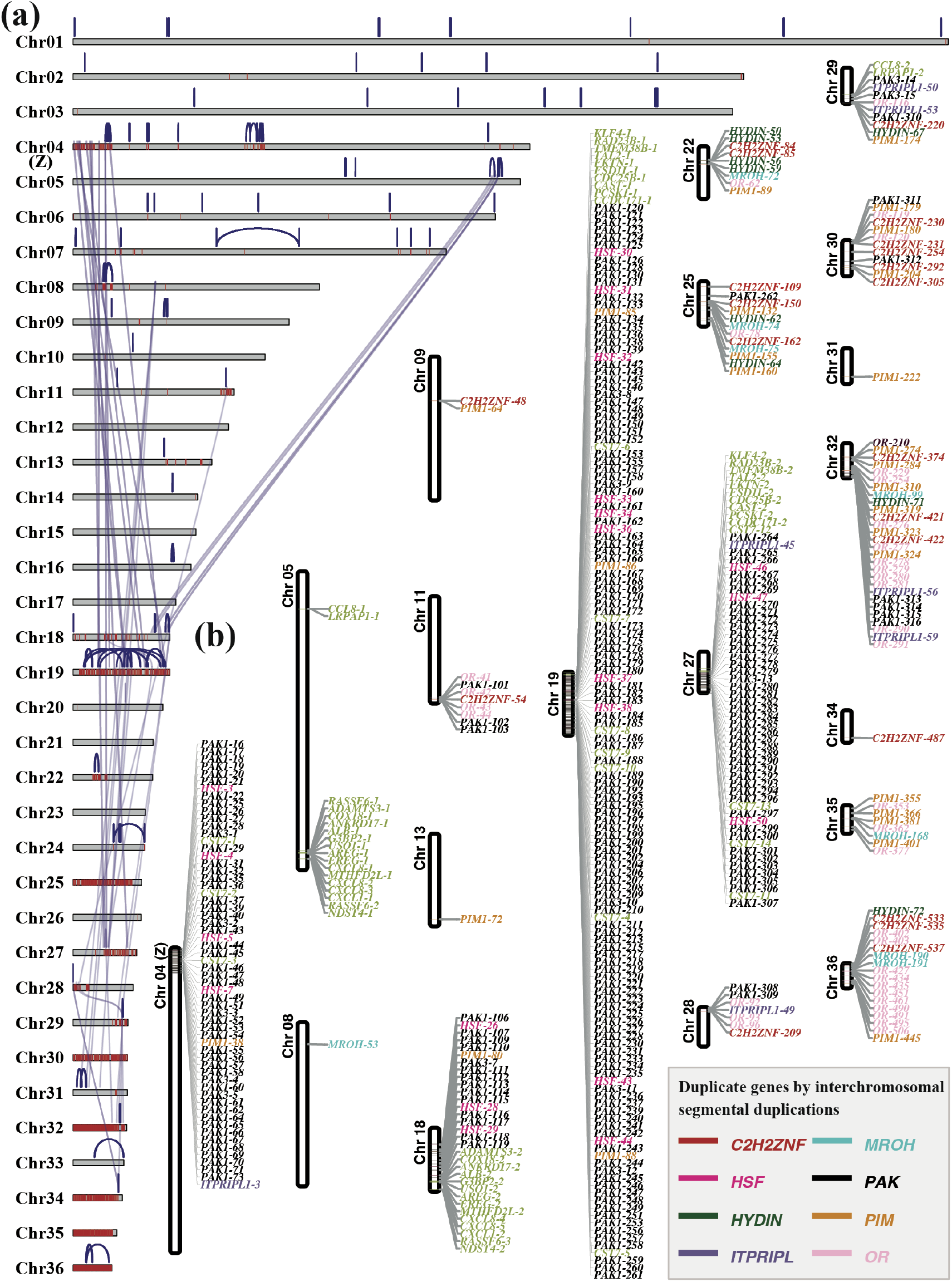
Segmental duplication contents in tree sparrow. (a) The pattern of segmental duplications of tree sparrow. The graphic shows an overview of large and high-identity intrachromosomal (blue) and interchromosomal (grey purple) segmental duplications (>70 kbp and >95% sequence identity). The red bar highlight regions represent the *C2H2ZNF, OR, PIM, PAK, MROH, HYDIN, HSF* and *ITPRIPL* gene families. (b) The chromosomal distributions of protein coding genes located in interchromosomal duplication regions.

**Figure 5.**
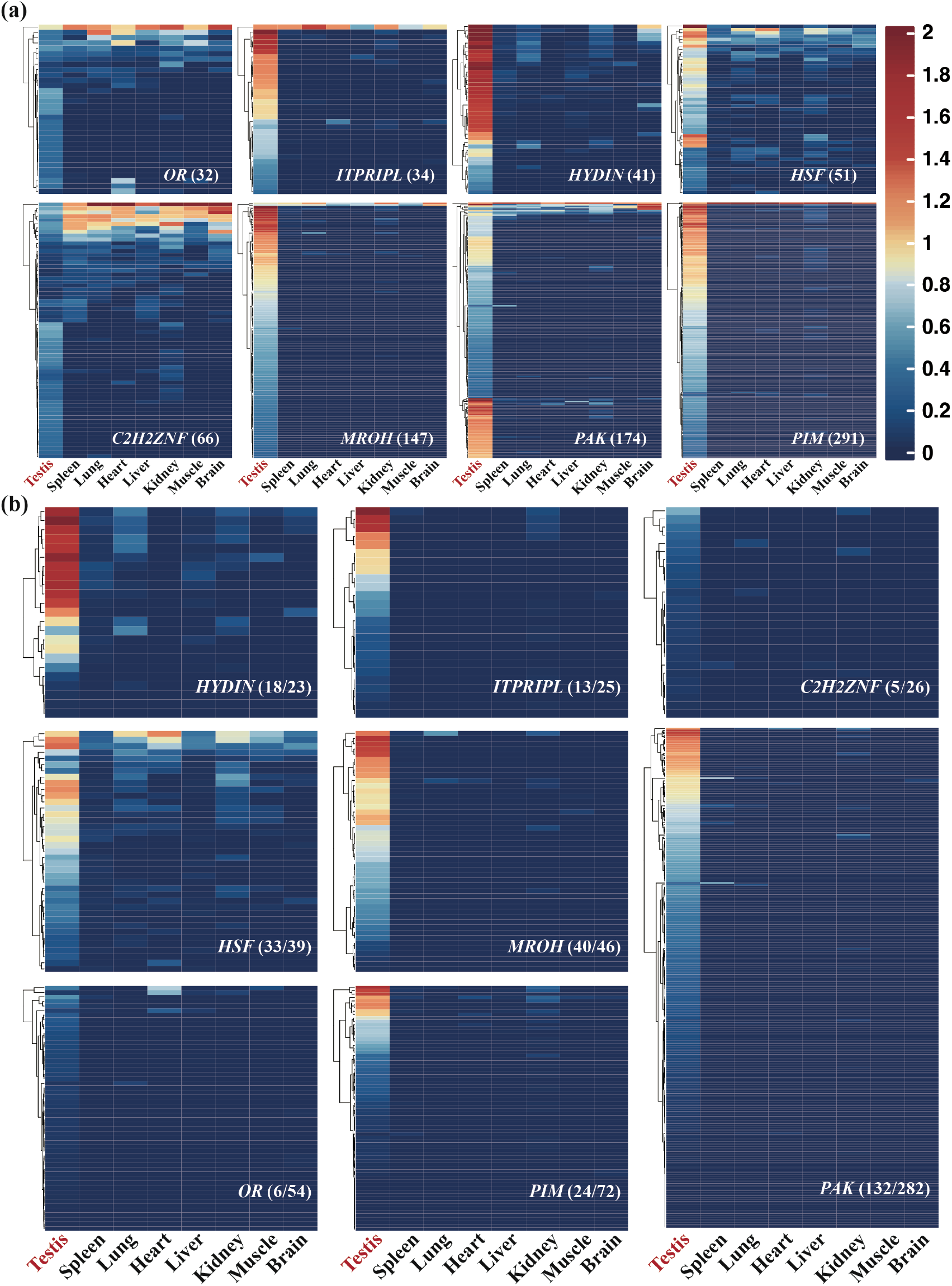
Tissue expression patterns of the eight significantly expanded gene families. (a) Heatmap of expression profiles of tree sparrow eight significantly expanded gene families. The transcriptionally inactive members were filtered and not shown in the heatmap, and the figures in brackets represent the numbers of transcriptionally active members. The scale bar represents the log_10_-transformed TMM values. (b) Heatmap of expression profiles of all SD genes, no matter transcriptionally active or inactive, of the eight families. The numbers in brackets represent active SD genes and all SD genes respectively.

### Landscape and comparative analysis of transposable elements

At least 18.27% of tree sparrow genome assembly is composed of repetitive elements, made up of transposable elements (TEs) (16.84%) and tandem repeat (1.43%) (Supplementary Table 4 and 5). The total TEs content is slightly higher than most of the 25 selected bird genomes, except for two species in Piciformes (*Picoides pubescens* and *Tricholaema leucomelas*) (Fig. 2c and Supplementary Table 6). DNA transposons compose 8.29% of the assembly and terminal inverted repeats (TIRs) elements account for most of the DNA transposons, whereas miniature inverted-repeat transposable elements (MITEs) and Helitrons only take up a small proportion (Supplementary Table 5). We noticed that the DNA transposons were clearly higher and showed greater expansion in tree sparrow than other birds (Fig. 2c), which were mainly derived from an ancient burst of TIR/DTC (CACTA) superfamily (Supplementary Fig. 6 and 7). Furthermore, a number of the DNA transposons are prevalent in passerines, indicating that they are potentially active in tree sparrow genome (Fig. 2d).

**Figure 6.**
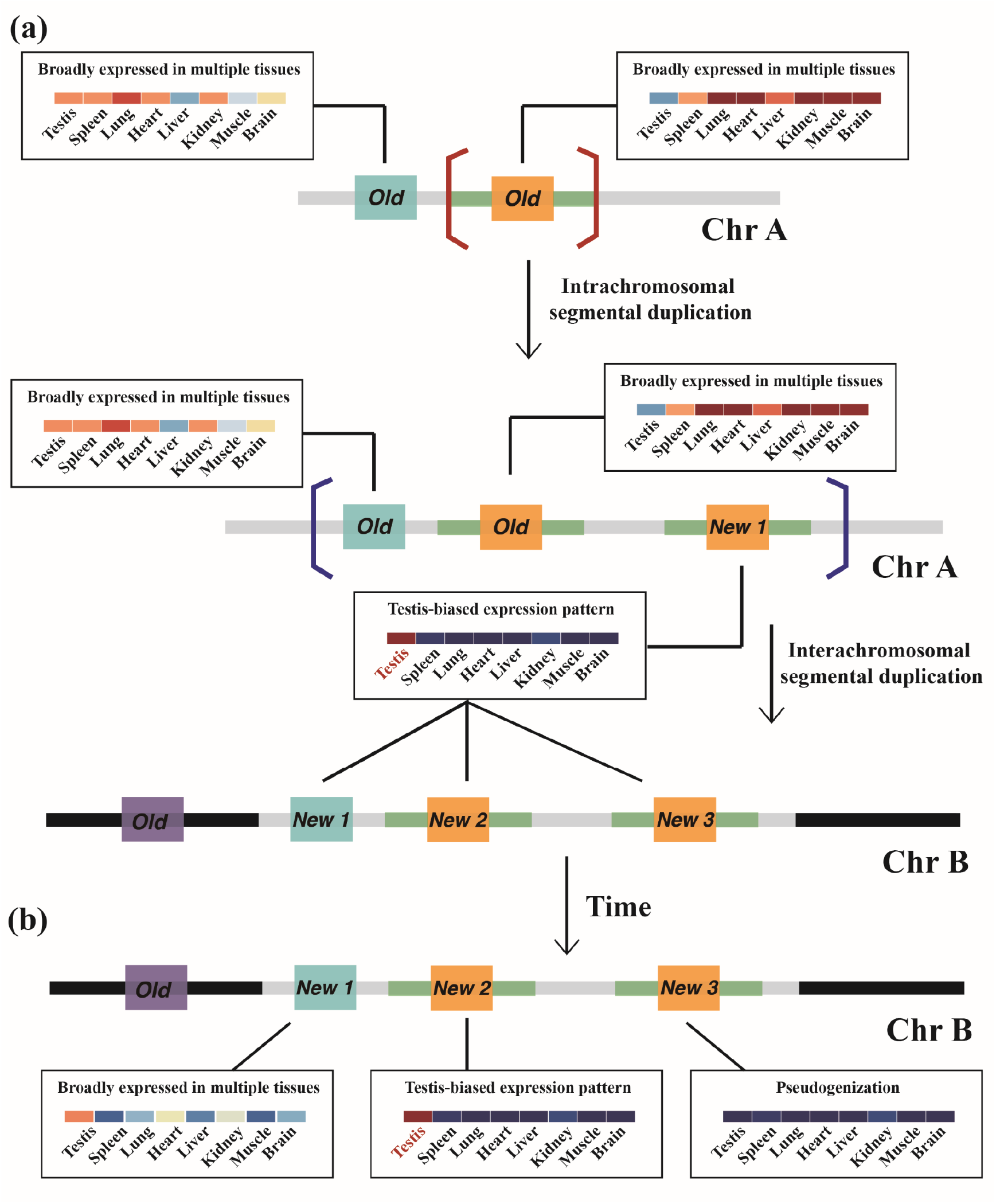
Model of SDs in tree sparrow. (a) Inter- and intrachromosomal duplication events occurred independently in tree sparrow genome. Following the SDs, the expression patterns of new genes were shifted to testis-biased. (b) Subsequently, the possible outcomes of new genes including becoming pseudogenes, maintaining testis-biased expression or changing to broadly expression in other tissues.

The long terminal repeat retrotransposons (LTR-RTs) are the most abundant retrotransposons in tree sparrow genome (Supplementary Table 5). The tree sparrow genome contains about 532 intact LTR-RTs, 442 of these elements are endogenous retroviruses (ERVs). The ERVs in tree sparrow genome were classified into 4 clades (betaretrovirus, gammaretrovirus, epsilonretrovirus and spumaretrovirus) using phylogenetic reconstruction of their reverse transcriptase (RT) domains (Fig. 2e). Betaretrovirus and gammaretrovirus are the two most common ERVs in tree sparrow and more betaretrovirus were detected in tree sparrow and zebra finch than in chicken genome (Fig. 2e). Furthermore, we found that a portion of tree sparrow and zebrafinch betaretrovirus RT domains were clustered with chicken alpharetrovirus (Fig. 2e).

Relative to LTR-RT, long interspersed elements (LINEs) and short interspersed elements (SINEs) are less common and active in tree sparrow genome as also in the other 5 songbirds (Fig. 2c and 2d). LINEs constitute about 3% of tree sparrow genome, when SINEs account for only 0.05% (Supplementary Table 5). CR1 elements are the domain LINEs in tree sparrow, but only a tiny fraction of them are potentially active (Supplementary Fig. 7).

The genomic landscape of transposable elements shows that the DNA transposons are relatively evenly distributed across chromosomes, accompanied by occasional scattered burst (Fig. 3), whereas the retrotransposons showed more complex and diverse distribution characteristics. For non-LTR retrotransposons, SINEs are rare in all chromosomes except for chromosome 9, when regions proximity to the assembled chromosomes termini often contain high density of LINEs (Fig. 3). Relative to large autosomes, LTR-RTs are more concentrated in Z chromosome and microchromosomes. Interestingly, we observed that LTR-RTs had the similar distribution trend with the eight significantly expanded gene families including *C*_*2*_*H*_*2*_*ZNF, OR, PIM, PAK, MROH, HYDIN, HSF* and *ITPRIPL* (Fig. 3).

### Segmental duplication contents and testis-biased expression pattern of new genes

Segmental duplications (SDs) are genomic sequences larger than 1 kbp that are duplicated at least one time in genome with high identity (>90%) (Bailey et al. 2001). In total, we identified 61.74 Mbp of nonredundant SDs (>1 kbp in length and >90% identity), which contained 692 annotated protein coding genes (Fig. 4a and Supplementary Table 8). Focusing on SD regions that carry genes, we detected expansions of 54 protein coding gene families through inter- and intrachromosomal duplications in tree sparrow genome (Fig. 4b and Supplementary Fig. 8). Among these families, *PAK* had the largest number of recently duplicated members (268 of *PAK1* and 14 of *PAK3*) and showed the most concentrated chromosomal distribution (Fig. 4b and Supplementary Fig. 8). In addition to *PAK*, the SD blocks also contained large numbers of copies (>20) of the other 7 significantly expanded gene families (*C2H2ZNF: 26, OR: 54, PIM: 72, MROH: 46, HYDIN: 23, HSF: 39*; *ITPRIPL: 25*) (Supplementary Table 8). However, members of each of these families showed relatively dispersed distribution patterns when compared with *PAK* (Fig. 4b and Supplementary Fig. 8).

Using the transcriptome data from different tissues (testis, spleen, lung, heart, liver, kidney, muscle and brain) of adult tree sparrow, we compared the expression profiles in different tissues of the eight significantly expanded gene families. Surprisingly, the highly transcribed genes from different families, whether located in the SD regions or not, generally exhibit testis-biased expression in 6 out of the 8 genes (Fig. 5). In contrast, the few members broadly expressed in different tissues are mainly located outside the SD blocks (Fig. 5). In addition, a large proportion of the members in these families, especially in *OR* (∼94%) and *C2H2ZNF* (∼89%), are almost not expressed in all tissues (Fig. 5a and Supplementary Fig. 9). The transcriptionally inactive genes are also common among SD genes (Fig. 5b).

Based on the above results, we inferred the pattern and process of SDs in tree sparrow (Fig. 6). For reasons has not yet been determined, bursts of both inter- and intrachromosomal duplication of several genomic regions occurred during the evolution of tree sparrow (Fig. 6a). Followed a series of SD events, a large number of additional new copies, mainly belonging to eight gene families including *C*_*2*_*H*_*2*_*ZNF, OR, PIM, PAK, MROH, HYDIN, HSF* and *ITPRIPL*, were added to tree sparrow genome. It seems that the expression status of new genes, no matter which families they belong to, were shifted to testis-biased expression pattern (Fig. 6a). Subsequently, a majority of new genes did not express in all tissues examined and became non-functional (pseudogenization), whereas some copies maintained the testis-biased expression or were expressed in other tissues (Fig. 6b).

## Discussion

Reference genomes are the cornerstone of modern genomics, and a high-quality assembly is valuable for providing insights into species evolution. We here assembled a chromosome-level genome of tree sparrow, which showed great improvement of both contig N50 (54.4 Mbp vs. 750.6 kbp) and scaffold N50 (64.7 Mbp vs. 11.1 Mbp) compared with a previous published short-read genome assembly based on short-read sequencing (Qu et al. 2020). The final assembly size of our assembly is larger than the previous one, which primarily caused by the increased assembled TE content (Supplementary Table 2). Due to the limitations of current NGS technology, just like the SDs, the estimates of TE content are always confused by highly repetitive region misassembly and collapse (Bailey & Eichler 2006; Bustos et al. 2016; Peona et al. 2018; Vollger et al. 2022). It seems that the TEs are underrepresented in the previous assembly of tree sparrow.

A great majority of bird genomes were previously reported to contain a low proportion of TEs (<15%), except for Piciformes (Feng et al. 2020). The TE content of tree sparrow (16.82%) is higher than most birds. However, unlike species in Piciformes, the higher TEs are derived mainly from expansions of DNA transposons and LTR-RTs (Fig. 2c), whereas the expansion of LINE type CR1 transposons contribute most for the higher level of TEs in Piciformes (Zhang et al. 2014; Manthey et al. 2018; Feng et al. 2020). As the scarcity of DNA transposons in avian genomes has been widely reported (Kapusta and Suh et al. 2016; Gao et al. 2017), we assumed that the unexpected expansion and recent activity of DNA transposons, especially CACTA superfamily, may be a species-specific or lineage-specific event in tree sparrow and may play an important role in genome evolution and speciation. We also noticed that most of intact LTR-RTs in tree sparrows are ERVs, which is common in birds (Bolisetty et al. 2012; Hayward et al. 2015; Kapusta and Suh 2016). There are some ERVs identified as betaretrovirus but more cluster with chicken alpharetrovirus, which may due to the evolutionary continuum leading from betaretroviruses to alpharetrovirus in birds (Bolisetty et al. 2012).

In addition to the minor expansion of TEs related to the other avian species, significant expansions of eight gene families including *C*_*2*_*H*_*2*_*ZNF, OR, PIM, PAK, MROH, HYDIN, HSF* and *ITPRIPL* were detected in the assembly. In addition, we noticed that these members from different families were always clustered together in chromosomes. This indicated that the expansion event of each family is not independent during evolution, while the different expansion scales of these families indicated the duplication also did not happene completely synchronously. Lots of members of these significantly expanded gene families were totally overlapped with the identified SDs blocks, and about 80% of the SD genes were members of the eight families, which suggested that inter- and intrachromosomal SDs caused a burst of new genes which are concentrated in the eight families. Duplicate genes are known as major sources of genetic material and evolutionary novelty, which play a crucial role in the adaptation to different environment (Moore and Purugganan 2003; Crow and Wagner 2006; Conant and Wolfe 2008; Magadum et al. 2013; Wang et al. 2022). The additional new copies added through SD may provide opportunities for tree sparrow adapting to new environments.

By analyzing the genomic region of the gene families which are related to the frequent and rapid SD events, we noticed that these eight gene families had similar chromosomal distribution pattern with LTR-RTs. On the one side, this result may indicate that an insertion site preference for LTR-RTs is exist in these families. Interestingly, the PIM, one of the eight families, have been known as a preferential proviral integration site for Moloney murine leukemia virus (Cuypers et al. 1984). On the other side, the adjacent distributions may also indicate that TEs were involved in the segmental duplication processes. The enrichments of TEs in SD regions were widely reported in mammals (Bailey 2001; Bailey et al. 2003; Cheung 2003; She et al. 2008) and insects (Fiston-Lavier et al. 2007; Zhao et al. 2013; Zhao et al. 2017), although the enriched TEs are different in different species. Despite all this, it still remained uncertain about whether the LTR-RTs mediated the SDs in tree sparrow, or some other mechanisms drove the duplication events and the expansion of LTR-RTs was just the by-products of SDs.

We then compared the transcription status of the significantly expanded gene families. Just as reported previously, pseudogenization is the most common fate of the duplicate genes (Lynch and Conery 2000), most of members of these families showed no expression in all tissues, even in the most recently duplicate copies (SD genes). In addition, among the transcriptionally active members, lots of testis-biased expressed genes were detected, and there still were some members showed broadly expressed pattern especially among the members outside the SD regions. Compared with the old genes, the new gene duplicates are more prone to have testis-biased or testis-specific expression, which have been verified in multiple species (Vinckenbosch et al. 2006; Cui et al. 2015; Kondo et al. 2017; Assis 2019; Zhang and Zhou 2019) and led to the “out of testis” hypothesis. This hypothesis posits that the promiscuous transcription in the testis and the powerful selection pressures such as sperm competition in the male germline encourage the emergence and fixation of new genes, and these new genes may be expressed and acquire new functions in other tissues later (Kaessmann 2010). The similar testis-biased expression pattern in eight gene families with diverse structure and functions in tree sparrow is consistent with the “out of testis” hypothesis in birds.

In conclusion, the high-quality chromosome-level assembly of tree sparrow improves our knowledge about the SDs in avian species. The SD events added a large number of new copies of eight gene families into tree sparrow genomes. These SDs and subsequent burst of new genes greatly shaped the tree sparrow genome and facilitated the evolutionary process. In addition, the testis-biased expression patterns of these new genes provide direct proof for the “out of testis” hypothesis. We hope that our study can inspire the further studies and exploration on the SDs and their evolutionary consequence in other avian species.

## Materials and methods

### Sampling and sequencing

All animal collections and experiments were approved by the Committee on the Ethics of Animal Experiments of School of Life Sciences of Lanzhou University. The muscle sample was obtained from a male tree sparrow caught by mist nets in 2021 from Liujiaxia (35°56′N, 103°53′E) of Gansu Province, China. DNA was extracted using the Qiagen DNeasy Blood and Tissue Kit. DNA concentration (minimum of 80 ng/µL) was measured using Qubit DNA Assay Kit in Qubit 2.0 Flurometer (Life Technologies, CA, USA). For PacBio sequencing, libraries were constructed with an average insert size of 15kb using SMRTbell Express Template Prep Kit 2.0 (Pacific Biosciences, Menlo Park, USA) and sequenced by PacBio Sequel II. Hi-C libraries were prepared following a standard protocol (Belton et al. 2012) and sequenced by Illumina HiSeq 4000 (Illumina, San Diego, USA). After filtering out low quality and duplicated reads, a total of 58.23 Gb (∼45 ×) of HiFi reads and 106.29 Gb (∼83 ×) of Hi-C reads were used for genome assembly.

### Genome assembly and annotation

Hifiasm version 0.16.0 (Cheng et al. 2021) was used for assembling PacBio HiFi reads into highly continuous and accurate contigs. HiC-Pro version 3.1.0 (Servant et al. 2015) was used to process Hi-C data from raw sequencing reads to normalized contact maps and the generated bin matrix results were taken as input data for EndHiC (Wang et al. 2021) to assemble hifiasm-assembled long contigs into chromosomal-level scaffolds.

RepeatModeler version 2.0.1 (Flynn et al. 2020) was used to construct a *de novo* repeat library for the assembled genome of tree sparrow. We employed RepeatMasker version 4.1.1 (Tarailo-Graovac and Chen 2009) to search for tandem elements by aligning the genome sequence against a combination of Repbase (Bao et al. 2015) database and the *de novo* repeat library constructed by RepeatModeler. Next, we used EDTA (Ou et al. 2019) pipeline to detect and annotate transposable elements (TE). Subsequently, the soft-masked genome was sent to MAKER version 3.01.03 (Holt and Yandell 2011) pipeline to predict protein-coding genes. All available protein sequences of zebra finch (*Taeniopygia guttata*), great tit (*Parus major*), house sparrow (*Passer domesticus*), European pied flycatcher (*Ficedula hypoleuca*), American crow (*Corvus brachyrhynchos*) and golden-collared manakin (*Manacus vitellinus*) from NCBI were aligned to the assembled genome using BLAST+ version 2.2.28 (Camacho et al. 2009) to provide protein homology evidence. All available RNA-seq reads of tree sparrow in public database were assembled into transcript using Trinity version 2.13.2 (Grabherr et al. 2011), and the transcript sequences were aligned to the genome to provide RNA evidence. After polishing those alignments around splice sites using Exonerate version 2.2.0 (Slater and Birney 2005), protein homology evidence and RNA evidence were integrated with *ab initio* gene predictions from SNAP (Korf 2004), AUGUSTUS version 3.4.0 (Stanke et al. 2008) and GeneMark-ES version 4.68 (Lomsadze et al. 2005) by MAKER. Finally, the functions of predicted gene sets were annotated by eggNOG-Mapper version 2.1.6 (Cantalapiedra et al. 2021). The accuracy and completeness of assembly and annotation were assessed by BUSCO version 5.2.2 (Manni et al. 2021).

### Synteny analysis and visualization of genomic landscape

We used MUMmer version 4.0.0 (Marçais et al. 2018) to align the entire assembly to the latest reference genome of chicken downloaded from Ensembl, and the syntenic dot plots of the whole genome and 15 longest assembled chromosomes were generated by web visualization tool Assemblytics (Nattestad and Schatz 2016). We performed pairwise complete CDS alignment among chicken, tree sparrow and zebra finch using MCscanX (Wang et al. 2012). The guanosine and cytosine (GC) content, gene density, TE density and tandem repeat density for each 500 kb genomic bin were calculated by BEDTools version 2.30.0 (Quinlan and Hall 2010) and shown in circular genome map by Circos version 0.69.8 (Krzywinski et al. 2009).

### Comparative genomic and phylogenetic analysis

Orthologous groups between tree sparrow and another 25 representative avian species, covering 13 orders (Accipitriformes, Anseriformes, Apterygiformes, Casuariiformes, Charadriiformes, Falconiformes, Galliformes, Passeriformes, Piciformes, Psittaciformes, Strigiformes, Struthioniformes, and Tinamiformes), were inferred using OrthoFinder version 2.5.4 (Emms and Kelly 2019). The obtained amino acid sequences of 4,085 one-to-one single copy orthologous proteins from the 26 species were aligned using MAFFT version 7.475 (Katoh and Standley 2013) and concatenated into a supergene. The concatenated alignment was used to construct a phylogenetic tree of 26 species using RAxML version 8.2.12 (Stamatakis 2014) with 100 bootstrap replicates. We ran MCMCtree program in PAML version 4.9 (Yang 2007) to estimate the species divergence time with two known divergence time points: between chicken and turkey (*Meleagris gallopavo*) (CI: 22-42 Mya) and between duck (*Anas platyrhynchos*) and swan goose (*Anser cygnoides*) (CI: 22-36 Mya) in TimeTree database (Kumar et al. 2022). We used CAFE version 4.2.1 (De Bie et al. 2006) to detect gene family expansion and contraction.

### TE analysis

We used the same EDTA pipeline as tree sparrow to annotate TE in other 25 bird genomes, in order to ensure comparability. Firstly, we used a combination of LTR_FINDER (Xu and Wang 2007) and LTRharvest (Ellinghaus et al. 2008) with LTR_retriever (Ou and Jiang 2018) to annotate LTR-RTs. We extracted the intact LTR-RTs to further classified using TEsorter (Zhang et al. 2022) with Gypsy Database (GyDB) (Llorens et al. 2011). The RT domains of the identified ERVs of tree sparrow, zebra finch and chicken were used to construct a maximum-likelihood (ML) tree using IQ-TREE version 2.1.2 (Minh et al. 2020). Secondly, we used the LINE and SINE repeat database in RepeatMasker to generate a library to annotate LINEs and SINEs. Finally, the DNA transposons were detected by TIR-Learner (Su et al. 2019) and HelitronScanner (Xiong et al. 2014). TIR-Learner was used to detect TIRs and MITEs, when HelitronScanner was used to detect Helitron transposons. TIRs and MITEs were classified into 5 different superfamilies: *hAT* (DTA), *CACTA* (DTC), *PIF/Harbinger* (DTH), *Mutator* (DTM), and *Tcl/Mariner* (DTT). We used the calcDivergenceFromAlign.pl script in RepeatMasker to calculate divergence rate using the Kimura 2-parameter divergence metric. Only TE with 0% divergence may be potentially active. The numbers of TEs and eight significantly expanded gene families (*C*_*2*_*H*_*2*_*ZNF, OR, PIM, PAK, MROH, HYDIN, HSF* and *ITPRIPL*) were counted in 1 Mbp windows with 200 kbp steps using BEDTools.

### Segmental duplication characterization

We used BISER version 1.2.3 (Išerić et al. 2022) to detect segmental duplication with identity >90% and length >1 kbp. The largest (>70 kbp) and most identical (>95%) segmental duplications were visualized using karyoploteR package (Gel and Serra 2017) in R. The protein coding genes overlapped with SDs blocks were extracted using BEDTools. The chromosome distributions of these genes were obtained from genome annotation information and visualized using TBtools version 1.098685 (Chen et al. 2020).

### Tissue expression profiles

We downloaded all valuable transcriptome data of tree sparrow from the NCBI Sequence Read Archive (SRA) database. The reads were mapped to the assembly using STAR v2.7.9a (Dobin et al. 2013). We performed gene-level quantification approach using featureCounts v2.8.1 (Liao et al. 2014) and the expression heatmaps of all members of eight significantly expanded gene families in eight tissues (brain, heart, kidney, liver, lung, muscle, spleen, and testis) were generated using ComplexHeatmap v2.10.0 (Gu et al. 2016) package in R.

## Data Accessibility Statement

All raw sequence data have been deposited in the National Center for Biotechnology Information (NCBI) Sequence Read Archive (SRA) (BioProject: PRJNA867105).

## Author contributions

Y. Z. and S.W. conceived the project and designed the research. S.W., Y.S., Z.L. and Y.M. collected samples in the field. S.W. performed the bioinformatic analysis and drafted the original manuscript. Y.Z., G.S. and Y.J. revised and edited the manuscript.

## Acknowledgments

This work was supported by the National Natural Science Foundation of China (Grant No. 31572216) and Foundation for Excellent Doctoral Student of Gansu Province (No. 22JR5RA413). We received support for the computational work from the Supercomputing Centre of Lanzhou University.

## Notes

### Competing Interest Statement

The authors have declared no competing interest.

